# Are Your Covariates Under Control? How Normalization Can Re-introduce Covariate Effects

**DOI:** 10.1101/137232

**Authors:** Oliver Pain, Frank Dudbridge, Angelica Ronald

**Affiliations:** Department of Psychological Sciences, Birkbeck, University of London, United Kingdom.; Department of Non-communicable Disease Epidemiology, London School of Hygiene and Tropical Medicine, United Kingdom.; Department of Health Sciences, University of Leicester, United Kingdom.

**Keywords:** Rank-based inverse normal transformation, normalization, covariates, confounding, residuals.

## Abstract

Many statistical tests rely on the assumption that the residuals of a model are normally distributed. Rank-based inverse normal transformation (INT) of the dependent variable is one of the most popular approaches to satisfy the normality assumption. Studies regularly adjust for covariates and then normalize the residuals. This study investigated the effect of regressing covariates against the dependent variable and then applying rank-based INT to the residuals. The correlation between the dependent variable and covariates at each stage of processing was assessed. An alternative approach was tested of applying rank-based INT to the dependent variable *before* regressing covariates was tested. Analyses based on both simulated and real data examples demonstrated that applying rank-based INT to the dependent variable residuals after regressing out covariates re-introduces a linear correlation between the dependent variable and covariates in almost all situations. This will increase type-1 errors and reduce power. Our proposed alternative approach, where rank-based INT was applied prior to controlling for covariate effects, gave residuals that were normally distributed and linearly uncorrelated with covariates. This approach is therefore recommended.

## Introduction

Many statistical tests rely on the assumption that the residuals of a model are normally distributed (Berry 1993). In genetic analyses of complex traits, the normality of residuals is largely determined by the normality of the dependent variable (phenotype) due to the very small effect size of individual genetic variants (Servin and Stephens 2007). However, many traits do not follow a normal distribution. In behavioral research in particular, questionnaire data often exhibit marked skew as well as a large number of ties between individuals. The non-normality of residuals can lead to heteroskedasticity (comparison of variables with unequal variance) potentially resulting in increased type I error rates and reduced power (Feingold 2002).

There are several approaches to either satisfy the normality assumption or control for violations of it. One of the most popular of these approaches is the transformation of the dependent variable to fit a normal distribution, i.e. normalization. There are several transformations that can be used for this purpose, the most popular being log, power or Box-Cox transformations, and rank-based inverse normal transformations (INTs), also referred to as quantile normalizations, such as the Van de Waerden transformation (Beasley et al. 2009). In many cases the use of log transformation has been shown to be insufficient in normalizing data. Conversely, rank-based INTs always creates a perfect normal distribution when there are no tied observations. Previous studies have reported that although rank-based INTs can lead to loss of information, this approach controls power and type-I error rate (Wang and Huang 2002; Peng et al. 2007). However, a comprehensive review of rank-based INTs demonstrated that in certain scenarios, rank-based INTs do not control type-I error, although they remain useful in large samples where alternative methods, such as resampling, are less practical (Beasley et al. 2009).

It is often desirable to adjust for covariates in analysis. In genetic studies, principal components of ancestry are commonly included to reduce confounding by population structure. When a transformation to normality is used, the covariates may be included in the analysis model after transformation, or alternatively they may be regressed against the response prior to the residuals being transformed to normality. The latter approach has been used in a number of recent high profile studies (Locke et al. 2015; Wain et al. 2015; Jones et al. 2016) and is also automated in the ‘rntransform’ function within GenABEL, a popular R package (Aulchenko et al. 2007). One reason is that in collaborative consortia, it is more convenient to perform in-house adjustments for covariates and transformations prior to data sharing. Another reason is that confounders may be considered to have their effects on the untransformed, rather than the normalized, variable. Finally, pre-adjustment for covariates will break many of the ties that are present in data derived from questionnaires or other rating scales that are usually represented by a small number of discrete values.

In this study we demonstrate that regressing covariate effects from the dependent variable creates a covariate-based rank, which is subsequently inflated by rank-based INT, leading to increased type-1 errors and reduced power. We use simulations to study the effects of regressing out covariates on quantitative data with and without ties, in terms of the degree of skew in the data and the proportion of tied observations. The effect of randomly splitting tied observations before rank-based INT will also be explored. Our results suggest that the practice of regressing out covariate effects prior to transformation should be discouraged. As an alternative we suggest that rank-based INT, randomly ranking tied observations, should be performed before controlling for covariates.

## Methods

### Simulation of phenotypic data

Two types of phenotypic data were simulated: quantitative variables containing no tied observations (herein referred to as continuous variables) and quantitative variables containing tied observations (herein referred to as questionnaire-type variables). These variables were simulated to exhibit different degrees of skew ranging from −2 to 2. Skewed variables were created using the R ‘rbeta’ function, which randomly generates numbers following a beta distribution with two shape parameters to control the degree of skew. Each simulated variable contained 10,000 observations. To create tied observations in the questionnaire-type variables, the initially continuous data were collapsed into evenly distributed and discrete response bins. The number of response bins, determining the proportion of tied observations, was varied between 5 and 160.

The R functions used to create continuous and questionnaire-type variables, called ‘SimCont’ and ‘SimQuest’ respectively, are available in Supplementary Text 1 and 2.

A normal distribution is defined by skew = 0 but also kurtosis = 0. Given that the simulated variables were generated to follow a beta distribution, variables with a skew equal to zero may not have a kurtosis equal to zero. To ensure that the correction of kurtosis was not driving effects seen when skew is equal to zero, continuous and questionnaire-type variables were also generated using the ‘rnorm’ function in R to exhibit both a skew and kurtosis of zero. The functions used to create continuous and questionnaire-type with skew and kurtosis fixed to zero, called ‘SimContNorm’ and ‘SimQuestNorm’ respectively, are available in Supplementary Text 3 and 4.

### Simulation of covariate data

To create correlated covariate data, noise was added to each simulated phenotypic variable until the desired phenotype-covariate correlation was achieved. Phenotype-covariate correlations (Pearsons) were varied between −0.5 and 0.5. Noise was added to the questionnaire variables using the ‘jitter’ function in R.

The R function used to create covariates for each phenotypic variable, called ‘CovarCreator’, is available in Supplementary Text 5.

### Testing the effect of rank-based inverse normal transformation after regressing out covariate effects

Linear regression of each covariate against the corresponding phenotypic variable was used to calculate phenotypic residuals, which are linearly uncorrelated with the covariates. The Spearman’s rank-based correlation between the phenotypic residuals and covariates was measured. Phenotypic residuals were then normalized using the‘rntransform’ from the GenABEL package in R, which applies a rank-based INT similar to van de Waerden transformation. To determine whether the transformed residuals were still linearly uncorrelated with covariates, the Pearson correlation between the transformed residuals and covariates was calculated.

### Testing the effect of applying rank-based inverse normal transformation (randomly splitting ties) before regressing out covariate effects

This was carried out using the same simulated questionnaire-type and continuous variables and covariates. The raw questionnaire-type and continuous variables underwent rank-based INT using a modified version of the ‘rntransform’ function from GENABEL that randomly ranks any tied observations. The modified version of ‘rntransform’, called ‘rntransform_random’, is available in Supplementary Text 6. Linear regression of each covariate against the corresponding normalized questionnaire-type and continuous variables was used to calculate phenotypic residuals, which are linearly uncorrelated with the covariates.

One concern with rank-based INT, particularly when randomly splitting ties, is that the linear relationship between the phenotypic variable and independent variables (including covariates) may be severely distorted. To determine the extent to which rank-based INT when randomly splitting ties distorts phenotypic variables, the Pearson correlations between the untransformed and transformed phenotypic variables were calculated. To determine the extent to which rank-based INT when randomly splitting ties distorts the relationship between the phenotypic variables and covariates, the Pearson correlation between the transformed phenotypic variables and covariates was calculated.

Another concern with normalizing the phenotypic variable before regressing out covariates is that the process of regressing out covariates may re-introduce skew in the residuals. To determine the extent to which regressing covariates from normalized phenotypic variables re-introduced skew, the skew of the residuals was assessed.

### Demonstration using real data

To determine whether the predicted effects (when using simulated data) of performing rank-based INT before or after regressing out covariate effects are accurate, the same procedure was applied to real questionnaire data provided by the Twins Early Development Study (TEDS)(Haworth et al. 2013). Data from two questionnaires were used measuring Paranoia and Anhedonia. Both of these measures are part of the SPEQ (Specific Psychotic Experiences Questionnaire)(Ronald et al. 2014). Individuals with missing phenotypic data were excluded from all analyses. Sum scores of unrelated individuals were calculated by summing the response of each item. Each item of both the Paranoia and Anhedonia scales were coded as values from 0-5, with the total ranges of the Paranoia and Anhedonia scales being 0-75 and 0-50 respectively. In order to make the real data comparable to simulated data, sum scores were calculated using different numbers of items (1, 2, 4, 8) to create the number of response bins similar to those in the simulations (5, 10, 20, 40). The covariates used were age (continuous variable skew of −0.32) and sex (binary variable with skew of 0.22). Table I shows the skew, number of response bins (proportion of ties) and correlation with covariates for each of dependent variable. The same analysis procedure for simulated data was carried out using the real TEDS data.

**Table I:** Skew, range, and correlation with covariates for dependent variables derived from TEDS sample.

| Dependent variable | Range | Skew | Pearson correlation with age | Pearson correlation with sex |
| --- | --- | --- | --- | --- |
| **Paranoia** | 5 | 1.357 | 0.055 | 0.018 |
| **Paranoia** | 10 | 1.195 | 0.043 | -0.026 |
| **Paranoia** | 20 | 1.095 | 0.030 | -0.022 |
| **Paranoia** | 40 | 1.296 | 0.022 | -0.059 |
| **Anhedonia** | 5 | 1.868 | -0.006 | 0.177 |
| **Anhedonia** | 10 | 0.858 | -0.025 | 0.127 |
| **Anhedonia** | 20 | 0.651 | -0.020 | 0.135 |
| **Anhedonia** | 40 | 0.537 | -0.013 | 0.205 |

## Results

### The effect of rank-based inverse normal transformation after regressing out covariate effects using simulated data

As expected, regressing covariates against phenotypic variables created phenotypic residuals that were linearly uncorrelated with covariates. Although there was no linear correlation, in almost all simulations a rank-based correlation existed between the residuals and covariates (Supplementary Figure 1-7). As a consequence, rank-based INT of residuals re-introduced a linear correlation between the phenotypic variables and covariates (Supplementary Figure 8-14). Three factors predicted to affect the extent to which rank-based INT of residuals re-introduced a correlation between the phenotypic variables and covariates were tested. These factors were the original skew of the phenotypic variable, the original correlation between the phenotypic variable and covariate, and the proportion of tied observations in the original phenotypic data.

First, in terms of skew, greater skew of the phenotypic variable was associated with a higher correlation between the normalized phenotypic residuals and the covariate data (Figure 1, Supplementary Figure 8-14). The direction of skew had no effect on the correlation between the normalized residuals and the covariate data. The effect of normalizing residuals when skew was equal to zero remained when kurtosis was also fixed to zero (Supplementary Figure 15).

**Figure 1.**
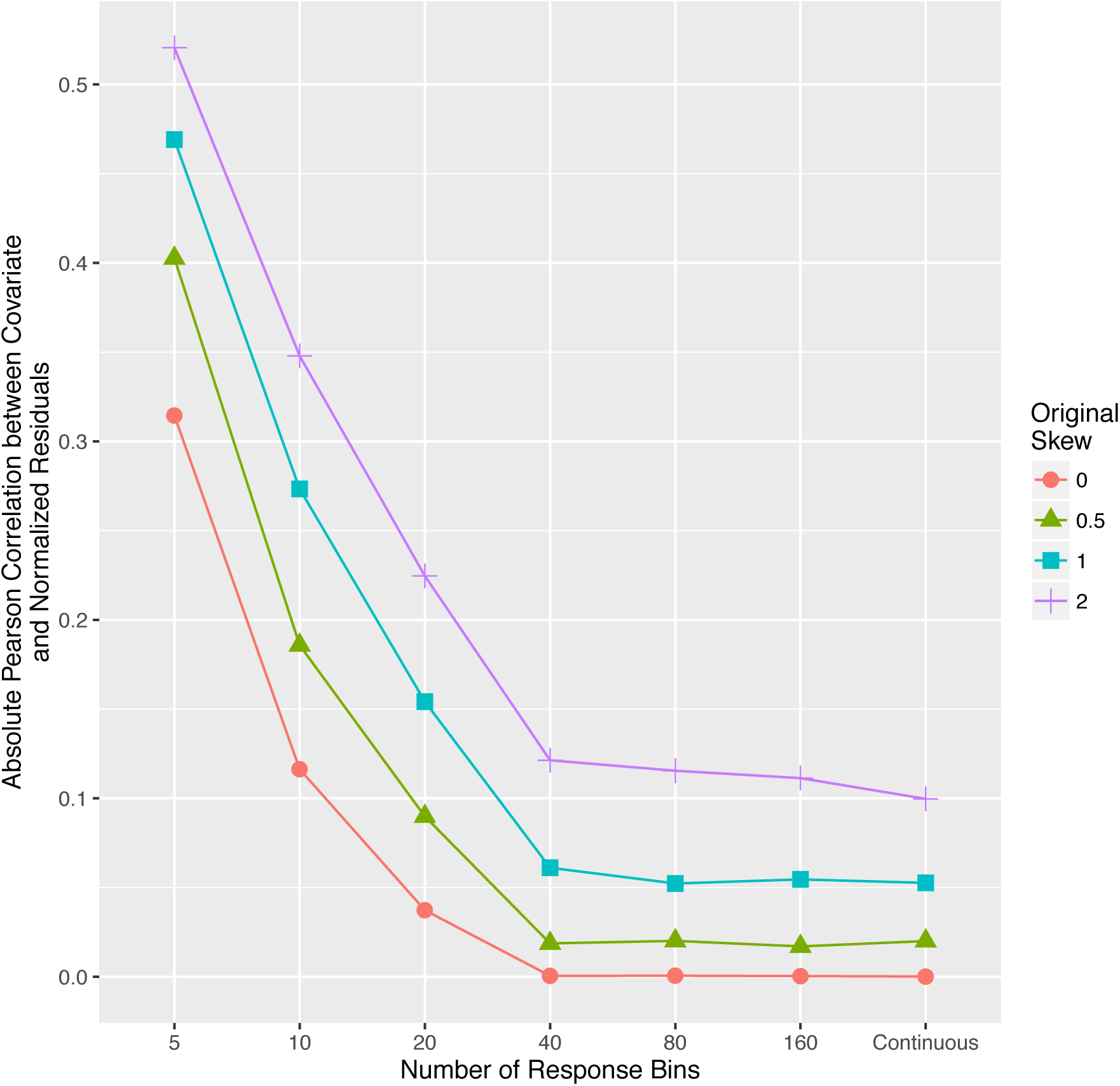
The relationship between the number of available responses (x-axis) and correlation between normalized residuals and covariate (y-axis) for different values of the skew in the raw phenotypic data. Within this figure, the correlation between the untransformed phenotypic data and covariate data is at 0.06.

Second, the direction of the original correlation between the original phenotypic variable and the covariates was reversed after rank-based INT of residuals. In questionnaire-type data, when the proportion of tied observations was high, the magnitude of correlation between the original questionnaire data and covariates had a negative relationship with the degree to which normalization re-introduced the correlation with covariates (Supplementary Figure 8). However, this negative relationship reversed as the proportion of ties decreased (Supplementary Figure 16). This means when the proportion of tied observations was low (or in continuous data), the magnitude of correlation between the original questionnaire data and covariates had a positive relationship with the degree to which normalization re-introduced the correlation with covariates.

Third, in terms of the proportion of ties in the phenotypic variable, a decreased number of response bins in the questionnaire-type data (i.e. smaller range and more tied observations) resulted in an increased correlation between covariates and normalized residuals (Figure 1). However, even when there were 160 response bins, or the data were continuous, rank-based INT still re-introduced a correlation with covariates when the data had an original skew >0.5 (Supplementary Figure 13 and 14).

As previously mentioned, although there is no linear correlation between phenotypic residuals and covariates, a rank-based correlation between the phenotypic residuals and covariates remained in almost all simulations. The factors affecting the magnitude of rank-based correlation between phenotypic residuals and covariates are the same as those influencing the effect of rank-based INT of residuals (Supplementary Figure 17-18).

### The effect of rank-based inverse normal transformation (randomly splitting ties) before regressing out covariates using simulated data

Rank-based INT of phenotypic variables, randomly splitting ties, before subsequent regression of covariates against the normalized phenotypic data, always resulted in phenotypic residuals with no linear correlation with covariates, and in the majority of simulations, skew less than 0.05.

The process of rank-based INT whilst randomly splitting ties decreased the correlation with covariates by a small amount (median change of 5%)(Supplementary Table 1). The extent to which the covariate correlation decreased was dependent on the original correlation between the covariate and the dependent variable, the skew of the dependent variable, and the proportion of tied responses in the dependent variable (Supplementary Figures 19-24).

The Pearson correlations between dependent variables before and after rank-based INT (randomly splitting tied observations) were between 0.77 and 1.00. An increased proportion of tied observations and increased skew led to a decreased correlation after rank-based INT (Supplementary Figure 25).

Regressing covariates after normalizing the dependent variables introduced a smaller degree of skew when covariates had either a low skew themselves or a low correlation with the dependent variable. The degree to which regressing covariate effects introduced skew was not dependent on the proportion of tied observations. Overall, regressing covariates introduced a small amount of skew to the dependent variable (0.00 – 0.11) unless the covariate had a correlation with the dependent variable over 0.25 and a skew greater than 0.05 (Supplementary Figure 26). However, highly skewed covariates may introduce larger amounts of skew even when exhibiting a low correlation with the dependent variable.

### Effect of rank-based inverse normal transformation of residuals when using real data

The observed effect of applying rank-based procedures to residuals within simulated questionnaire-type data was validated using real questionnaire data from TEDS. When using the age covariate (continuous) the magnitude and direction of effect of applying rank-based procedure to residuals were similar to those of simulated questionnaire-type data (Supplementary Table 2-3). The effect of rank-based procedures on residuals when using real questionnaire data was slightly reduced in comparison to effects observed when using simulated questionnaire-type data.

When the sex covariate (binary) was used, the magnitude, and in some cases the direction, of the effect of rank-based procedures varied from effects observed in simulated data. Although regressing the effect of a binary covariate altered the outcome of rank-based procedures, application of rank-based procedures to residuals still re-introduced a correlation with covariates (Supplementary Table 4-5). Importantly, when a dichotomous variable was used, a large number of ties in the data still existed reducing the efficacy of rank-based INT.

### Effect of rank-based inverse normal transformation before regressing out covariates when using real data

The effect of rank-based INT (randomly splitting tied observations) before regressing out covariate effects in real questionnaire data was comparable to the effects observed when using simulated data. Rank-based INT, randomly splitting ties, and subsequent regression of covariates created residuals that were linearly uncorrelated with covariates and normally distributed (Supplementary Table 6-7). The correlation between the dependent variable and covariate did vary slightly before and after rank-based INT (Supplementary Table 6-7). Contrary to the observed effects when using simulated data, the correlation between the dependent variable and the covariate did not always decrease. The Pearson correlation between raw and normalized questionnaire data varied between 0.83 and 0.99 dependent on the skew of the raw data and the number of response bins (Supplementary Table 8). Similar to the results of our simulations, the effect of regressing covariates out of the normalized variables did not re-introduce skew greater than 0.02 in any situations (Supplementary Table 6-7).

## Discussion

This study has demonstrated that regressing covariates against the dependent (phenotypic) variable and then transforming the resulting residuals to normality re-introduces a correlation between the covariates and the normalized dependent variable. This effect occurs because the process of regressing covariates against the response variable leads to a covariate-based rank in the residuals, which is then used to redistribute the data (Figure 2). This effect of regressing covariates against response variables occurs when the response variable is continuous (contains no tied observations) or questionnaire-type (contains tied observations), however the effect increases as the proportion of tied observations increases. The degree to which the covariate correlation is re-introduced during rank-based INT is dependent on the original skew of the response variable, although when the data contain a large proportion of tied observations, a correlation with covariates is re-introduced even when there is no skew.

**Figure 2.**
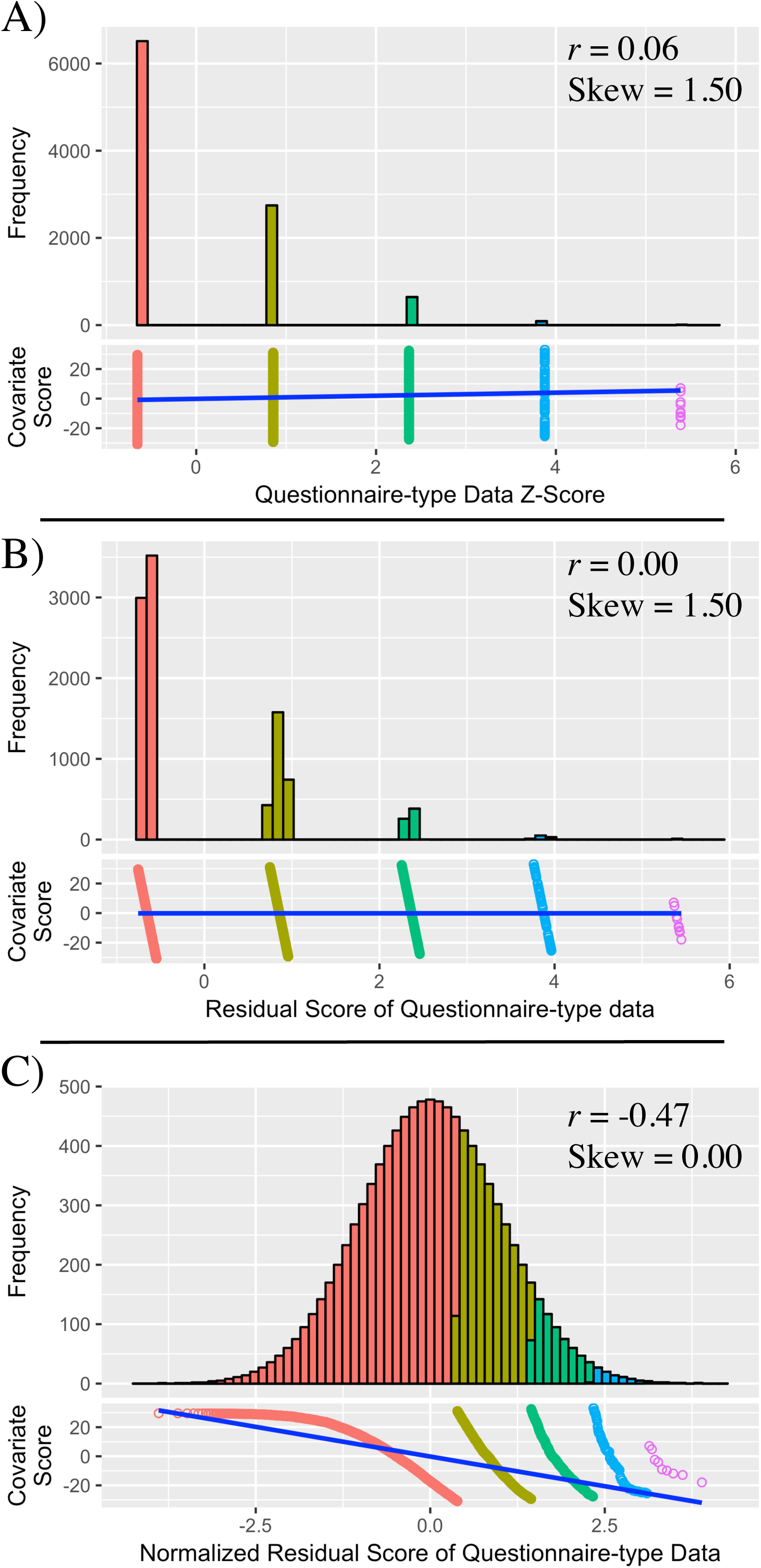
The effect of applying a rank-based INT to residuals of questionnaire-type data, i.e. after regressing out covariates. All correlations referred to in this figure are Pearson (linear) correlations. A) Untransformed questionnaire-type variable and its relationship with a continuous covariate. The questionnaire-type variable has a range of 5. A weak linear relationship exists between the questionnaire-type variable and covariate. B) Questionnaire-type variable residuals after regressing out the relationship with the covariate. No linear relationship exists between the questionnaire-type residuals and covariate. Regressing out covariate effects has led to the separation of many tied observations, creating a covariate-based rank within the questionnaire-type variable residuals. C) After the rank-based INT of questionnaire-type variable residuals, the transformed questionnaire-type variable residuals show a strong linear correlation with the covariate. This correlation is stronger and in the opposite direction to the original correlation between the untransformed questionnaire-type variable and the covariate.

This study has also evaluated an alternative procedure for preparing data for parametric analyses, whereby the response variable undergoes rank-based INT, randomly separating ties, before regressing out covariate effects. Our findings demonstrate that this alternative approach is preferable as it creates a normally distributed response variable with no correlation with covariates (Figure 3). The notion of normalizing the response variables before estimating its relationship with covariates may seem counterintuitive as the process of normalization may disrupt the true relationship between variables. Although this may be true in some scenarios, when the variables are skewed and/or contain tied observations, the change in relationship between variables due to normalization (Supplementary Table 6-7) is small relative to the change in relationship when normalizing residuals (Supplementary Table 2 and 4). In contrast, regressing covariates after normalization will leave no correlation between the variables, meaning that any confounding by those covariates will be eliminated.

**Figure 3.**
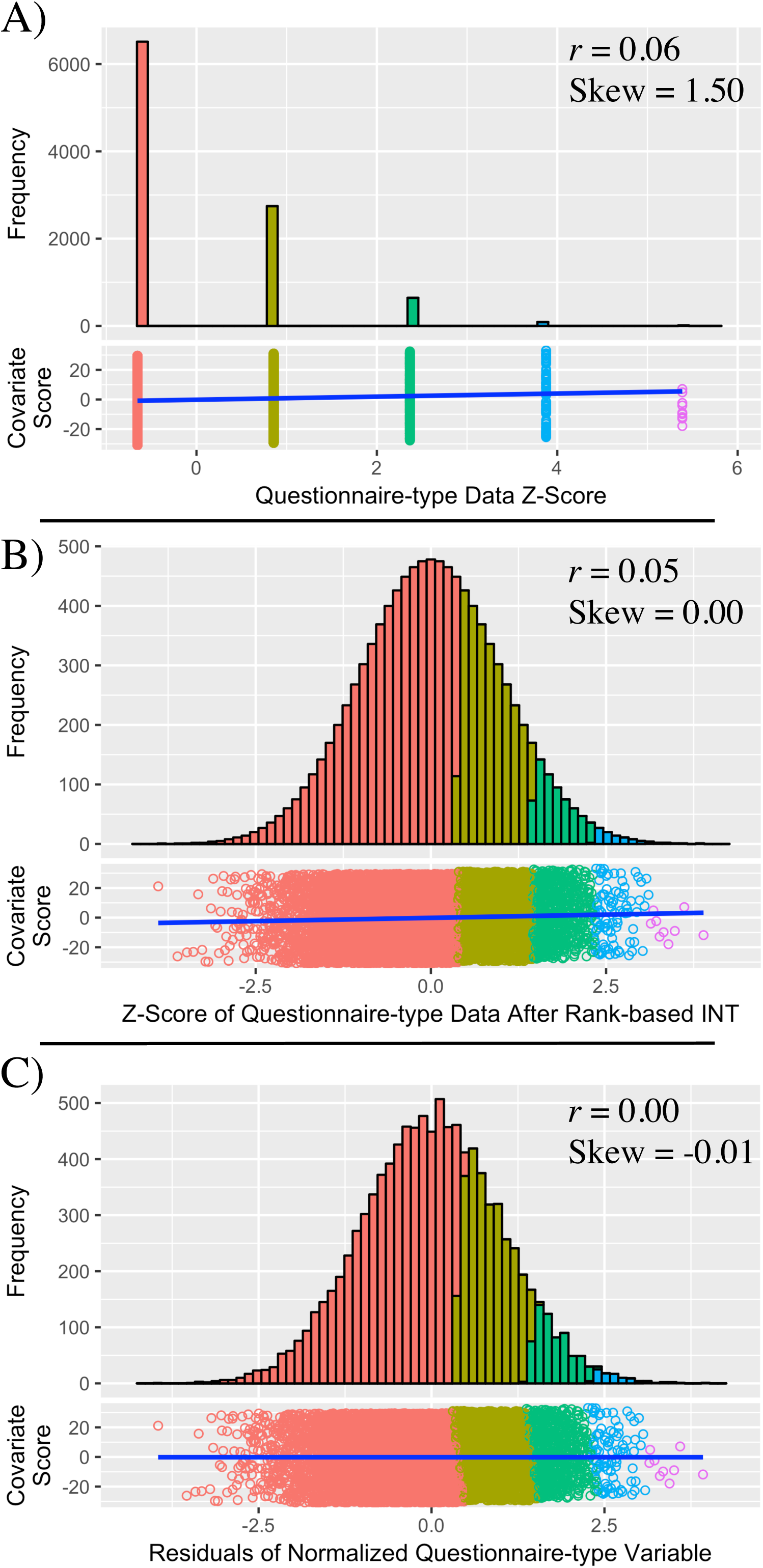
The effect of applying a rank-based INT to questionnaire-type data before regressing out covariates. All correlations referred to in this figure are Pearson (linear) correlations. A) Untransformed questionnaire-type variable and its relationship with a continuous covariate. The questionnaire-type variable has a range of 5. A weak linear relationship exists between the questionnaire-type variable and covariate. B) Questionnaire-type variable after rank-based INT, randomly splitting tied observations. Relationship between the questionnaire-type variable remains intact. C) Covariate effects have been regressed from the normalized questionnaire-type variable. There is no linear relationship between the residuals and the covariate, and the skew is close to zero.

Given the importance of phenotypic transformations, authors must describe the details of this process. Many studies do not clearly describe the details in which the data is processed, however there are some studies that have clearly applied rank-based INT to residuals (Locke et al. 2015; Wain et al. 2015; Jones et al. 2016). We do not believe that the results of these studies are seriously in error as they have either dealt with traits that have a very low skew and/or are continuous, or they have replicated their findings using binary outcomes based on untransformed data. However, we do believe that the potential problems with rank based INT of residuals after correcting for covariates are not well known, and that researchers should be aware of these issues before applying such a procedure. We suggest that, when possible, researchers adjust for covariates after rather than before applying a normalizing transformation, or employ other methods that do not assume normality.

These findings are not just relevant to rank-based INT of residuals but highlight the importance of procedures that introduce or alter the rank of observations. Another procedure that will alter the rank-based relationship between variables is the calculation of factor scores via principal components analysis (PCA). PCA is a method that applies orthogonal transformation to identify linearly uncorrelated axes of covariance among observations. Although the derived factors are linearly uncorrelated, they may have a rank-based correlation. Therefore, if the factors are skewed, subsequent rank-based INT will introduce a linear correlation between factors. Similar to the example of normalizing residuals, if the original correlation between the latent variables is positive, rank-based INT will lead to a negative correlation between derived factors.

Beyond demonstrating that rank-based INT of residuals introduces a confounding correlation with covariates, this study has also provided evidence that applying other rank-based methods, such as non-parametric, to residuals may also lead to confounding.

Although, this study concludes that normalization of the dependent variable should be performed prior to the regression of covariates, regressing out covariates that are either highly skewed or highly correlated with the dependent variable, may introduce substantial skew to the residuals. However, this scenario is considered unlikely.

## Conclusion

This study has demonstrated that rank-based INT of phenotypic residuals after adjusting for covariates can lead to an overcorrection of covariate effects leading to a correlation in the opposite direction between the normalized phenotypic residuals and covariates, and in questionnaire-type data, often of a greater magnitude. This finding has implications for all rank-based procedures and highlights the importance of clearly documenting how the raw data is handled. Normalization of phenotypic data before regressing out covariates has been explored as an alternative procedure and has been shown to produce normally distributed phenotypic residuals that are uncorrelated with covariates.

## Acknowledgments

We thank Doug Speed for providing helpful comments on the manuscript prior to submission. We also thank Robert Plomin, Andrew McMillan, the TEDS research team, and their participants for providing TEDS data for use in this study.

## Compliance with ethical standards

### Ethical approval

All procedures performed in studies involving human participants were in accordance with the ethical standards of the institutional and/or national research committee and with the 1964 Helsinki declaration and its later amendments or comparable ethical standards. This article does not contain any studies with animals performed by any of the authors.

### Informed Consent

Informed consent was obtained from all individual participants included in the study.

